# Function and interactions of a protein bridge between the inner membrane complex and subpellicular microtubules in *Toxoplasma gondii*

**DOI:** 10.1101/2024.05.29.595897

**Authors:** Emily S. Cheng, Andy S. Moon, William D. Barshop, James A. Wohlschlegel, Peter J. Bradley

## Abstract

*Toxoplasma gondii* is an obligate intracellular parasite that utilizes peripheral membrane and cytoskeletal structures for critical functions such as host cell invasion, replication, and maintaining cellular morphology. These structures include the inner membrane complex (IMC) as well as the underlying longitudinal subpellicular microtubules (SPMTs) that provide support for the IMC and give the parasite its distinctive crescent shape. While the IMC and SPMTs have been studied on their own, the mechanisms linking these adjacent structures remain largely unknown. This study identifies a *T. gondii* protein named IMT1 that localizes to the maternal IMC and SPMTs and thus appears to tether the IMC to the microtubules. We disrupt the IMT1 gene to assess function and then use deletion analyses and mutagenesis to reveal regions of the protein that are necessary for binding to the IMC cytoskeleton or SPMTs. Using proximity labelling with IMT1 as bait, we identify a series of candidate interactors in the IMC or SPMTs. Exploration of two of these candidates reveals that IMT1 regulates the levels of the microtubule associated protein TLAP2 and binds directly to the cytoskeletal IMC protein IMC1. Taken together, these interactions unveil the specific interactions linking two key cytoskeletal structures of the parasite and provides new insight into the organization of the structural backbone of *T. gondii*.

## Introduction

*Toxoplasma gondii* is an obligate intracellular parasite that infects every mammal, including ∼30% of the world’s human population (1). While infection in healthy individuals is typically asymptomatic, infections in immunocompromised individuals and in primary infections of pregnant women cause serious and even fatal disease (2). *T. gondii* is also a model system for related parasites in the phylum Apicomplexa, including *Plasmodium spp*., which cause malaria, and *Cryptosporidium spp.*, which cause diarrheal diseases in children (3). The establishment and maintenance of the intracellular niche of these parasites within their mammalian host cells is dependent upon parasite shape, which plays important roles in gliding motility, host cell invasion, and intracellular replication (4–6).

In *T. gondii* tachyzoites, the parasite exists as its hallmark crescent shape that is approximately 2uM in width by 6uM in length (7, 8). The pointed apical end of the parasite hosts the microtubule-based conoid, which serves as the site of secretion of secretory organelles involved in motility and host cell invasion (9, 10). The shape of the body of the parasite is dependent on several unique cytoskeletal and membrane structures at the periphery of the cell. One of these structures is the pellicle, which is the plasma membrane plus an underlying cytoskeletal and membrane organelle called the inner membrane complex (IMC) (8, 11). The IMC consists of flattened membrane vesicles called alveoli that are organized into a series of rectangular plates directly beneath the plasma membrane. The alveoli are supported by a second component of the IMC known at the IMC network, which is an array of intermediate filaments organized into an interlaced structure that provides support for the alveolar membrane vesicles (12, 13). The pellicle is further supported by 22 subpellicular microtubules (SPMTs) which originate at the apical polar ring at the base of the conoid and extend in a spiral pattern about two third the length of the cell (14). Together, the pellicle and SPMTs maintain the parasites shape and rigidity that enable efficient motility, host cell invasion, and the ability to adapt to the diverse environmental conditions encountered during its lifecycle.

The distinct layers of the pellicle and the underlying SPMTs are closely opposed to one another at the periphery of the parasite. They layers are tethered together by protein constituents which aids in association of the subcompartments and is important for function. For example, the glideosome motor complex that drives parasite motility associates with both the cytoplasmic face of the plasma membrane and the membranes of the alveolar vesicles (15, 16). The glideosome complex is anchored in the IMC membranes via the transmembrane domain of GAP50 and interacts with transmembrane micronemal adhesins on the plasma membrane. In addition, the GAP50 binding partner GAP45 is tethered to both the IMC and plasma membranes via acylation. Similarly, the IMC membranes are linked to the underlying cytoskeletal IMC network. Many of the intermediate filament proteins called alveolins that make up the IMC network are predicted to be palmitoylated, enabling association of the cytoskeleton with the IMC membranes (12, 13). The underlying SPMTs are regularly spaced and are closely associated with the IMC network, indicating a tight association of these components of the cytoskeleton (17, 18). While the precise mechanism is unclear, the IMC localized GAPM proteins serve to stabilize the SPMTs, indicating an interaction of the SMPTs and the IMC (19).

The 22 SPMTs underlying the IMC are remarkably stable and able to resist depolymerization by detergent extraction or cold treatment (20). The stability of the SPMTs is believed to be mediated by an array of microtubule associated proteins (MAPs) that may also provide a means for interaction with the IMC network. The first SPMT MAPs identified were SPM1 and SPM2 of which SPM1 contains six 32 amino acid repeats that function in SPMT binding (21). While SPM2 is dispensable, SPM1 plays a role in parasite fitness and is necessary for maintaining the stability of the SPMTs during detergent extraction. Subsequently identified MAPs include the thioredoxin-like proteins TrxL1 and TrxL2 and a series of four of TrxL1 associating proteins (TLAPs 1-4) (22, 23). Binding of TrxL1 to the microtubules is dependent on SPM1, and only TrxL2 and TLAP2 are able to directly bind microtubules when expressed in human cells. While individual knockouts often show minor no or minor phenotypes, a Δ*tlap2*Δ*spm1*Δ*tlap3* triple knockout results in significant growth and microtubule stability defects (22). While many components of the IMC and SPMT have been identified and investigated, the precise interactions that bridge these peripheral cytoskeletal structures remain poorly understood.

We and others have used proximity labelling to identify many of the protein constituents of both the IMC membranes and network (5, 24–27). In this study, we characterize the protein TGGT1_248740, which we previously identified and appears to have a novel localization to both the IMC and the SPMTs. We assess localization using widefield as well as Ultrastructure-expanded Microscopy (U-ExM) and find that the protein localizes to the IMC and SPMTs in maternal parasites, but not in developing daughter buds. We also assess function by gene knockout in both type I and type II strains and then use deletion analyses to identify regions of the protein that are important for targeting to the SPMTs or IMC. To identify candidate interacting partners of IMT1, we use proximity labeling which points to TLAP2 as a potential interactor on the microtubules and IMC1 as a partner in the IMC. Examination of TLAP2 in the Δ*imt1* parasites showed a substantial decrease of the protein on the SPMTs, suggesting that IMT1 stabilizes TLAP2 on the microtubules. In addition, we demonstrate that the minimal IMC binding region of IMT1 binds directly to the IMC1 by pairwise yeast two-hybrid (Y2H) analyses. Together, this work identifies a protein bridge between the microtubules and the IMC, dissects functional regions of the protein, and identifies specific interactions linking the SPMTs to the IMC.

## Results

### TGGT1_248740 localizes to the maternal IMC and SPMTs

Our previous proximity labelling experiments targeting using IMC proteins as bait have yielded an array of novel candidate IMC proteins that were endogenously epitope-tagged to determine their specific localizations (5, 24–26, 28). These studies resulted in the identification of many proteins that localize to distinct membrane or cytoskeletal subregions of the organelle including the IMC body, apical cap, and IMC sutures. One interesting protein we identified was TGGT1_248740, which we observed to localize to the subpellicular microtubules underlying the IMC but has not been explored further since its identification (5).

TGGT1_248740 has a predicted mass of 140.1 kDa and appears to be restricted to Toxoplasma and other members of the coccidia (29, 30). Although the protein lacks identifiable functional or structural domains, its N-terminus contains a multitude of lysine and glutamate-enriched charged repeats (e.g. KEEK), while its C-terminus has a series of eight tandem repeats of ∼15 amino acids containing the sequence “GYYDENG” and three predicted palmitoylation sites at residues 997, 998, and 1077 (Fig 1A, Fig S1) (31). The protein has a basic PI of 8.52, a feature seen in other microtubule binding proteins in both *T. gondii* and other systems that is believed to facilitate binding to the acidic tubulin dimer (21, 32).

**Figure 1.**
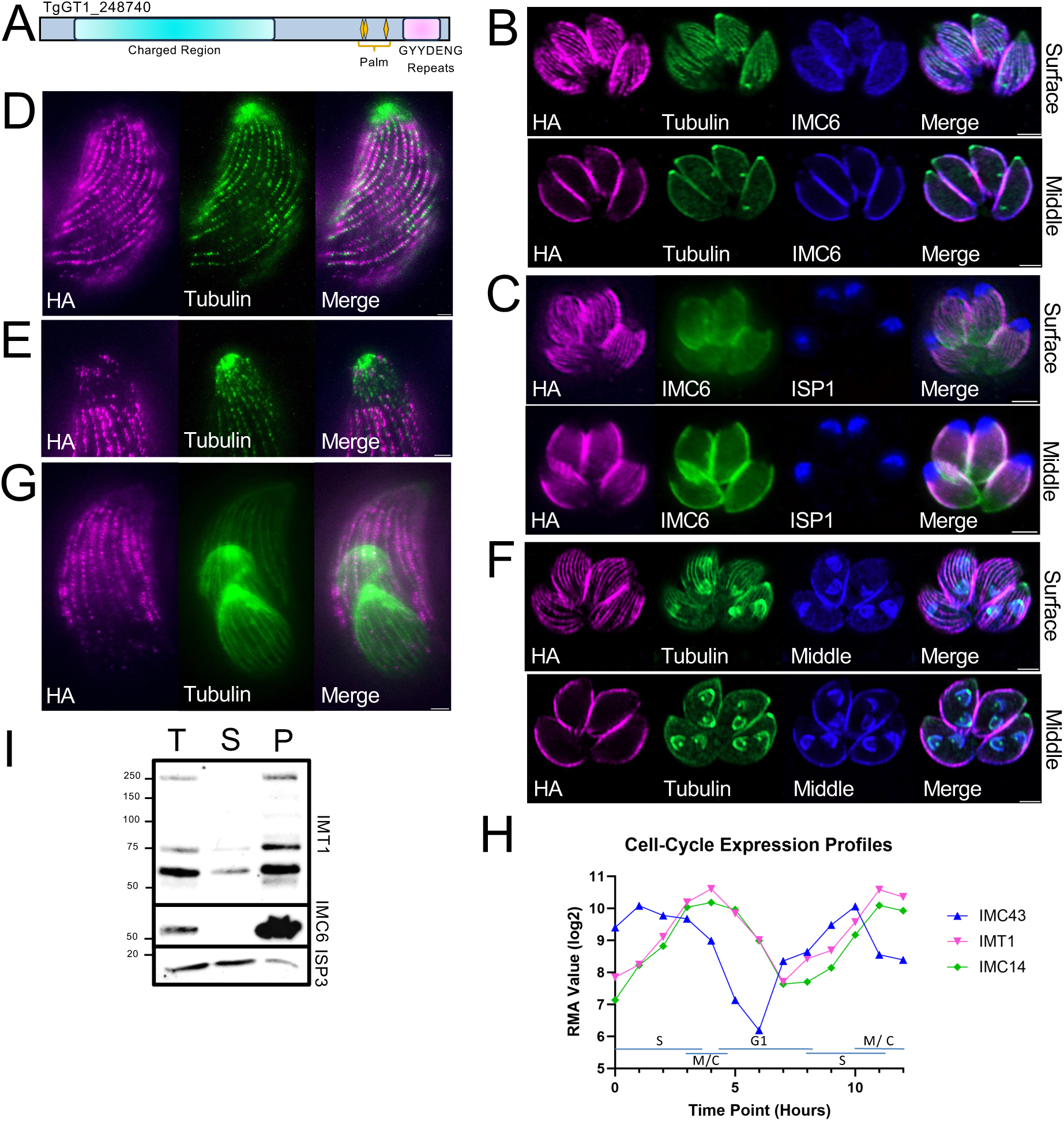
TGGT1_248740 localizes to the maternal IMC and SPMTs. A) Diagram of the protein sequence of TGGT1_248740 showing the position of a charged N-terminal region, C-terminal tandem repeats, and predicted potential palmitoylation sites. B) IFA of TGGT1_248740^3xHA^ focusing on the SPMTs at the surface of the parasite and colocalized with tubulin (top panel) as well as colocalized with IMC6 focusing on the IMC in a central plane of the vacuole highlighting peripheral localization (bottom panel); Magenta: mouse anti-HA; Green: eGFP-tubulin; Blue: rabbit anti-IMC6. C) IFA showing TGGT1_248740^3xHA^ is not present in the apical cap as assessed by ISP1 staining. Blue: mouse anti-ISP1; Green: rabbit anti-IMC6; Magenta: rat anti-HA. D) U-ExM of TGGT1_248740^3xHA^ showing a series of parallel lines running down the parasite body colocalized with tubulin highlighting the SPMTs. Magenta: rabbit anti-HA; Green: mouse anti-α-tubulin. E) U-ExM focused on the apical end of the parasite showing TGGT1_248740^3xHA^ is largely absent from the apical cap. Magenta: anti-HA; Green: mouse anti α-tubulin. F) IFA of parasites undergoing endodyogeny showing TGGT1_248740^3xHA^ is only detected in the maternal parasite and is not present in developing daughter buds; Magenta: mouse anti-HA; Green: eGFP-tubulin; Blue: rabbit anti-IMC6. G) U-ExM of parasite undergoing endodyogeny showing TGGT1_248740^3xHA^ is restricted to the maternal parasite. Magenta: rabbit anti-HA; Green: mouse anti-α- tubulin. H) The cell-cycle expression profile for TGGT1_248740 closely aligns with the cyclical pattern of the known maternal IMC protein IMC14, and expression peaks later than the early daughter bud protein IMC43. RMA = robust multi-array average. I) Western blot analysis of TX-100 detergent fractionation shows TGGT1_2248740^3xHA^ partitions to the cytoskeletal pellet with the IMC6 and ISP3 serving as cytoskeletal and membranous fraction controls, respectively. T = total, S = detergent soluble supernatant, P = detergent insoluble cytoskeletal pellet. All scale bars = 2 μm.

We reexamined the localization of TGGT1_248740 to assess its localization more carefully to the SPMTs and IMC using endogenously epitope tagged parasites (TGGT1_248740^3xHA^). Visualization of the SPMTs and IMC is challenging to capture in a singular plane as the SPMTs form a corset that encompass the circumference of the parasite, making them the most visible in surface views of the parasite, while the IMC exhibits the clearest peripheral staining in middle sections (22). Thus, we imaged parasites in two different focal planes to capture both structures (Fig 1B). When focusing on the microtubules at the surface, we confirmed SPMT localization by colocalization with GFP- tubulin engineered into the TGGT1_248740^3xHA^ tagged strain (Fig 1B) (33). When focusing on the central plane of TGGT1_248740^3xHA^ parasites, we saw that the protein colocalized well with the cytoskeletal IMC marker IMC6. While this could reflect the microtubules in this section, we reasoned that it might also localize to the IMC and serve as a bridge between the two structures at the periphery of the parasite.

We noticed that TGGT1_248740 staining was largely absent from the apical end of the parasite corresponding to the apical cap. Co-localization with the apical cap marker ISP1confirmed that TGGT1_248740 localizes to the body of the IMC, but not the apical cap (Fig 1C) (34). We also used U-ExM to examine the localization of TGGT1_248740 in further detail, which confirmed the protein localizes to the SPMTs and is mostly absent from the apical cap (Fig 1D, E) (35). We additionally found by both widefield and U-ExM that TGGT1_248740 staining is entirely absent from the daughter buds during endodyogeny, indicating this protein is restricted to the maternal SPMTs and IMC (Fig 1F, G). This agrees with cell cycle transcriptomic data which shows that TGGT1_248740 expression is cyclical with peak expression closely mirroring that of the maternal restricted IMC protein IMC14 and occurring later than the early IMC daughter bud protein IMC43 (Fig 1H) (36). Given the apparent localization of TGGT1_248740 to both the IMC and the SPMT, we named the protein IMC and SPMT protein 1 (IMT1).

To determine whether IMT1 is tethered to the parasite’s cytoskeletal elements, we performed a detergent extraction experiment which separates the detergent insoluble microtubules and IMC network from the detergent soluble membrane components, soluble proteins, and material loosely associated with the cytoskeleton (24). While IMT1^3xHA^ suffered significant breakdown during sample processing, it clearly partitioned to the insoluble cytoskeletal fraction with the IMC network component IMC6 and was not released from the membranes by detergent extraction like the palmitoylated membrane protein ISP3 (Fig 1I). This data demonstrates that IMT1 is firmly associated with some component(s) of the parasite’s cytoskeleton.

### Disruption of IMT1 does not affect parasite growth in vitro or cyst burden in vivo

IMT1 was assigned a phenotype score of –4.02 in the genome-wide CRISPR/Cas9 screen (GWCS), suggesting that it is either important or essential for parasite fitness (37). To directly assess IMT1 function, we disrupted IMT1 using CRISPR/Cas9 in IMT1^3xHA^ parasites. We selected a clonal isolate that lacked staining for the HA tag and confirmed the knockout by PCR (Δ*imt1*, Figs 2A-B). We then complemented the Δ*imt1* parasites by adding a full-length copy of IMT1^3xHA^ driven from its endogenous promoter and targeted to the UPRT locus which restored IMT1 staining in the SPMTs and IMC (IMT1^comp^, Fig 2C, D) (38). The levels of IMT1 expression in the IMT1^comp^ strain were confirmed to be similar to that of the IMT1^3XHA^ strain by western blot analysis (Fig 2E). Despite its strongly negative phenotype score, the Δ*imt1* strain showed no apparent morphological defects, gross effects on the microtubules, or significant growth defects in plaque area (Fig 2F-H).

**Figure 2.**
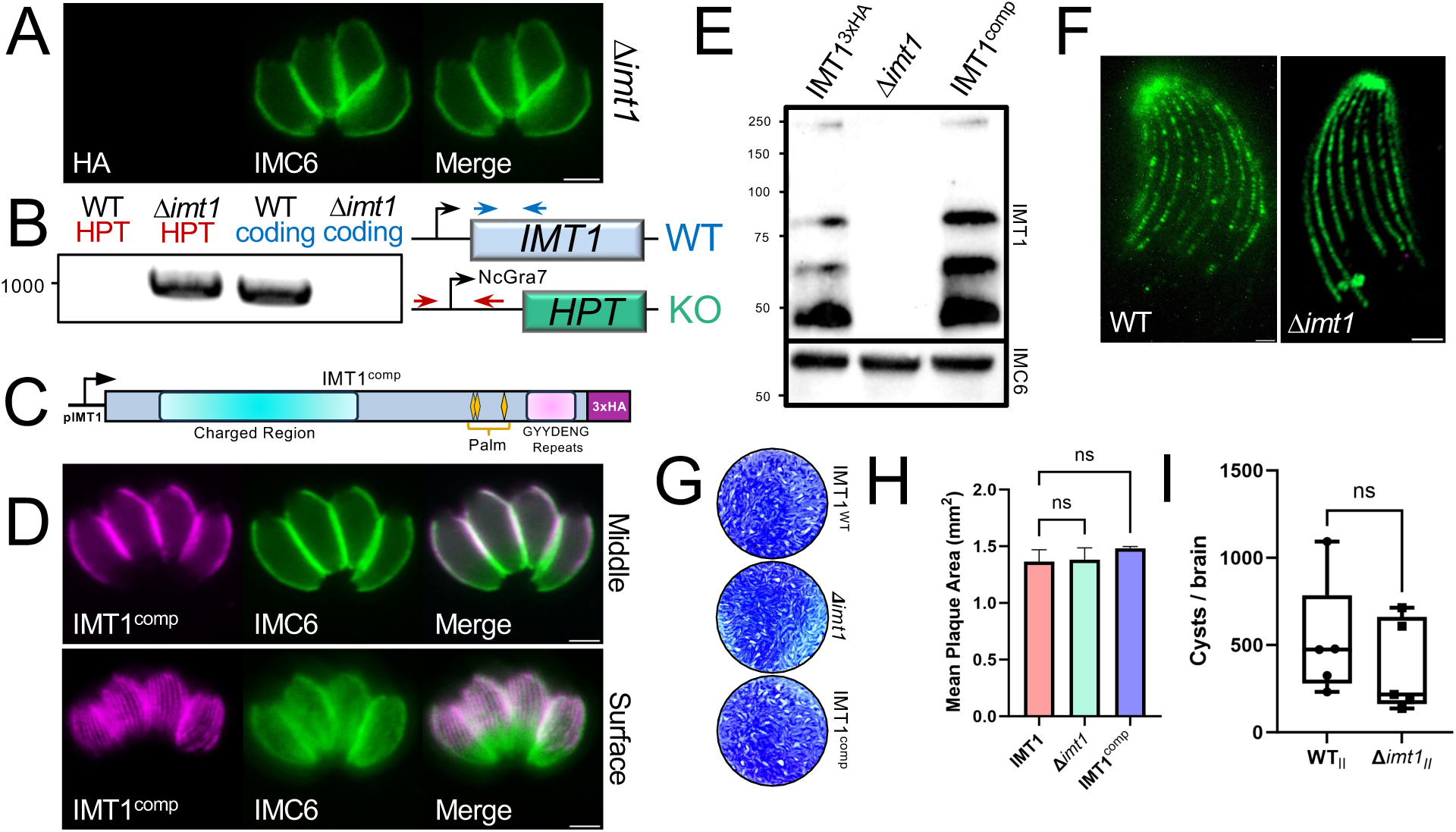
Gene knockout of IMT1. A) IFA of Δ*imt1* parasites lacking staining for the HA epitope tag. Magenta: mouse anti-HA; Green: rabbit anti-IMC6. B) PCR and diagram showing the Δ*imt1* strain contains the correct amplicon for the integrated selectable marker hypoxanthine-xanthine-guanine phosphoribosyl transferase (HPT) cassette and lacks the replaced IMT1 coding amplicon. The converse is true for the wildtype control, and all amplified bands match the sizes predicted by the primer positions shown in the diagram. C) Diagram of the IMT1 complementation construct driven from its own promoter. The construct is driven to the UPRT locus by UPRT flanking regions (not shown). D) IFA of IMT1^comp^ restores the IMC and SPMT localization seen in IMT1^3xHA^ parasites. The top panel focuses on the IMC in a medial view; the bottom panel focuses on the SPMT at the surface of the parasite. Magenta: mouse anti-HA; Green: rabbit anti-IMC6. E) Western blot of IMT1^3xHA^, Δ*imt1* and IMT1^comp^ parasites shows comparable expression levels of IMT1 protein in HA-tagged and complemented stains, but no HA-staining in the knockout. IMC6 is used as a loading control. F) U-ExM of WT and *Δimt1* show no gross difference in microtubule number or morphology. Green: mouse anti-α-tubulin. G) Images of representative wells after 7-day plaque assays of IMT1^3xHA^, Δ*imt1* and IMT1^comp^ strains. H) Quantification of plaque assays shows no significant difference in average plaque area between the three strains, indicating no lytic cycle defect is caused by the *IMT1* knockout. I) Brain cyst burden of CBA/J mice infected with type II parental control and Δ*imt1^II^* measured 30 dpi. Statistical analysis performed via unpaired t-test and reveals no significant difference between mice infected with the parental PruΔ*hxgprt*Δ*ku80* (WTII) or Δ*imt1II* as shown in box plots with min and max whiskers. All scale bars = 2 μm.

The disparity between the fitness score and the phenotype of our knockout could be the result of compensation during the knockout process, although we were unable to find any clear paralogues of IMT1 that might be able to compensate for its function. Using BLAST analysis we discovered the protein TGME49_316635 that has very low sequence similarity to IMT1 (4e^-6^), but contains charged repeats in the C-terminal region of the protein. TGME49_316635 appears to be only expressed in the sexual stages of the parasite and epitope tagging in both wildtype and Δ*imt1* parasites did not reveal any detectable expression in tachyzoites, thus it was excluded as a potential compensatory protein (Fig S3). We also tried to conditionally knockdown IMT1 using both auxin-inducible degron and tet-regulatable promoter replacement strategies but the degron resulted in incomplete knockdown and the promoter replacement failed to regulate expression.

Transcriptional analyses of the *T. gondii* acute versus chronic infection in mice indicates that IMT1 is upregulated in the chronic infection (Fig S2A) (39). To determine if IMT1 plays a role in cyst production or maintenance, we tagged and disrupted the gene in type II Prugniaud strain parasites that have been engineered to express *ldh2*-GFP specifically in the bradyzoite stage of the parasite (IMT1II, Δ*imt1*II, Fig S2B-E) (40). Similar to the type I knockout, Δ*imt1*II parasites showed no defect in vitro by plaque assays (Fig S2F, G). We then infected mice with parental and Δ*imt1*II parasites and evaluated the infection for 30 days post infection before assessing cyst burden (Fig. S2H). While a slight trend towards lower cyst numbers was present, it was not statistically significant (Fig. 2I). We therefore conclude that IMT1 is dispensable for both in vitro growth and for in vivo cyst burden.

### The C-terminus of IMT1 is necessary for binding to the IMC and SPMT

To investigate the localization of IMT1 to both the IMC and SPMTs, we assessed which regions of IMT1 could be responsible for binding to either component of the parasite’s cytoskeleton. As the N-terminus of IMT1 is rich in lysine and glutamate residues that are reminiscent of microtubule binding domains, we predicted that the N-terminus would be important for binding to the SPMTs (21, 32), while the C-terminus might bind to the IMC. To test this, we first constructed a series of five deletions from the C-terminus of IMT1^3xHA^, guided by predicted secondary structure analysis (Fig 3A). Each of deletions plus a full-length control (IMT1^wt^) were targeted to the UPRT locus in wildtype parasites (38). The localization of IMT1^wt^ matched that of endogenously tagged IMT1, confirming proper targeting and demonstrating that expression of a second copy of the protein does not affect localization of the protein (Fig 3B). Deletion of the C-terminal repeat domain in the IMT1^Δ1116-1283^ construct resulted in peripheral staining that colocalized with IMC6, indicating that this region is dispensable for localization to the periphery (Fig 3C). The IMT1^Δ1116-1283^ protein also localized to the SPMTs, although a small amount of the protein also appeared to be mislocalized from the microtubules (Fig 3C, bottom panel). Further analysis of expanded IMT1^Δ1116-1283^ parasites via U-ExM also showed colocalization with the microtubules in the parasite body, albeit with significantly greater cytoplasmic signal compared to endogenously tagged IMT1 (Fig 3D, Fig 1D). In spite of slightly dispersed staining in the body of the parasite, IMT1^Δ1116-1283^ was absent from the apical cap, matching wildtype localization (Fig 3E). All of the other deletions resulted in a loss of wildtype localization and either mislocalized to the cytoplasm or cytoplasm plus nucleus (Fig. 3F- I). This data demonstrates that the C-terminal region of IMT1 adjacent to the repeats is necessary for IMT1 localization and that the C-terminal repeats play a minor role in IMT1 SPMT localization.

**Figure 3.**
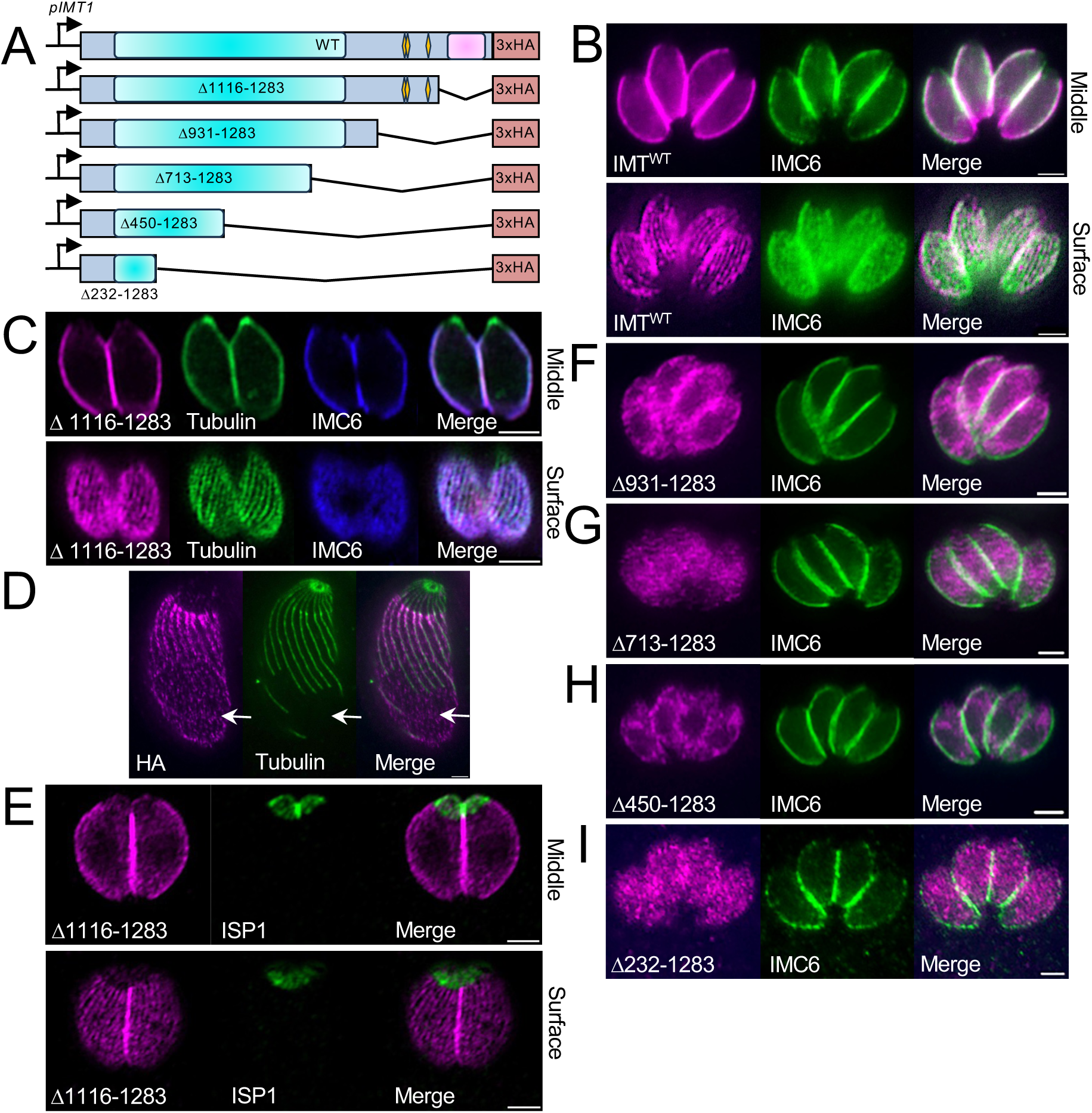
Deletion of the C-terminus mislocalizes IMT1. A) Gene model of the full- length second copy of IMT1 and five C-terminal deletions all driven from the endogenous promoter and fused to a 3xHA-epitope tag. All constructs contain flanking regions for targeting to the UPRT locus. B) IFA of the full-length second copy of IMT1 shows localization which matches that of the endogenously tagged line. Top panel focused on IMC; Bottom panel focused on SPMT. Magenta: mouse anti-HA; Green: rabbit anti-IMC6. C) IFA with top panel focused on the medial plane showing IMT1^Δ1116-1283^ properly localizes to the IMC. Bottom panel is focused on the surface of the parasite and reveals co- localization with tubulin. Magenta: mouse anti-HA; Green: eGFP-Tubulin; Blue: rabbit anti-IMC6. D) U-ExM of IMT1^Δ1116-1283^ shows co-localization with tubulin and dispersed staining throughout the parasite body indicating localization to the SPMT with slight mistargeting (arrows). Magenta: rabbit anti-HA; Green: mouse anti α-tubulin. E) Co- staining of IMT1^Δ1116-1283^ with ISP1 did not overlap, indicating IMT1^Δ1116-1283^ localizes to the IMC body but not the apical cap. Top panel: middle plane, Bottom panel: surface plane. Magenta: rabbit anti-HA; Green: mouse anti-ISP1. F-I) IFAs of the remaining four C- terminal localizations which mislocalize to the cytoplasm (IMT1^Δ931-1283^, IMT1^Δ450-1283^), the nucleus (IMT1^Δ713-1283^), or both (IMT1^Δ223-1283^). Magenta: mouse anti-HA; Green: rabbit anti-IMC6. All scale bars = 2 μm.

### The N-terminus of IMT1 is important for binding to the SPMTs

We similarly constructed a series of four N-terminal deletions to identify regions of IMT1 that are important for targeting (Fig 4A). IFA analysis of the IMT1^Δ2-449^ mutant showed peripheral staining that colocalized with IMC6 and also colocalized with the microtubules in surface staining (Fig 4B). However, examination of the microtubules by U-ExM revealed some mislocalization off the microtubules in the deletion mutant (Fig 4C). In addition, we noticed some of the IMT1^Δ2-449^ deletion protein localized to the apical cap (Fig. 4C, D). The IMT1^Δ2-712^ mutant localized to the periphery with IMC6, but appeared to largely lose microtubule localization (Fig 4E, F). The IMT1^Δ2-712^ mutant also showed some localization to the apical cap (Fig 4G). This loss of microtubule staining and presence in the apical cap was confirmed by U-ExM (Fig 4F). The IMT1^Δ2-930^ mutant showed a similar localization to the IMT1^Δ2-712^ mutant, but we also observed localization at the basal end of the parasite (Fig 4H-J). The largest deletion, IMT1^Δ2-1115^, resulted in the protein completely mislocalizing and largely residing in the parasite’s nucleus (Fig 4K).

**Figure 4.**
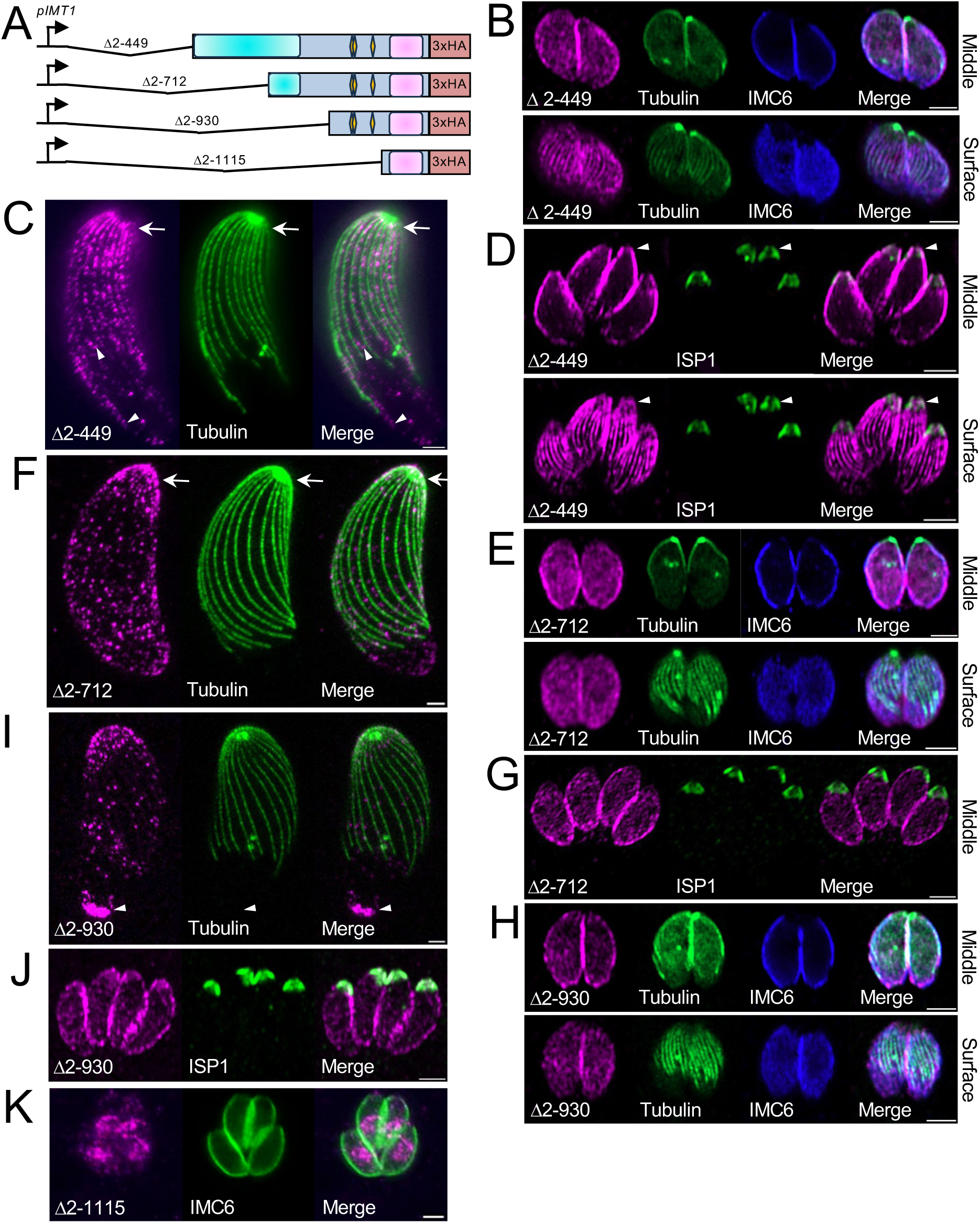
N-terminal deletion analyses identify regions responsible for SPMT and IMC targeting. A) Gene model of four N-terminal deletions driven from the endogenous promoter and fused to a 3xHA-epitope. All constructs are targeted to the UPRT locus via flanking regions that are not depicted. B) IFA of IMT1^Δ2-449^ focused on the parasite periphery shows colocalization with IMC6 and tubulin in middle and surface images, respectively. Magenta: mouse anti-HA; Green: eGFP-Tubulin; Blue: rabbit anti-IMC6. C) U-ExM of IMT1^Δ2-449^ co-localizes with tubulin and reveals staining at the apical end of the parasite not found with wildtype IMT1 (arrows); Magenta: rabbit anti-HA; Green: mouse anti-α-tubulin. D) IMT1^Δ2-449^ localized with ISP1 in both middle (top panel) and surface (bottom panel) views shows partial localization to the apical cap (arrowheads). Magenta: rabbit anti-HA; Green: mouse anti-ISP1. E) IMT1^Δ2-712^ also colocalizes with IMC6 in a middle view, but fails to localize with tubulin in a surface view. Magenta: mouse anti-HA; Green: eGFP-Tubulin; Blue: rabbit anti-IMC6. F) U-ExM of IMT1^Δ2-712^ shows disperse staining in the body of parasite that does not co-localize with tubulin, but is present in the apical cap (arrows) and displays increased staining at the basal end of the parasite; Magenta: rabbit anti-HA; Green: mouse anti-α-tubulin. G) Like IMT1^Δ2-449^, IMT1^Δ2-712^ colocalizes with ISP1 and loses exclusion from the apical cap. Magenta: rabbit anti-HA; Green: mouse anti-ISP1. H) IMT1^Δ2-930^ co-stains with IMC6 when focused on the periphery, but fails to co-stain with tubulin when focused on the parasite’s surface; Magenta: mouse anti-HA; Green: eGFP-Tubulin; Blue: rabbit anti-IMC6. I) U-ExM of IMT1^Δ2-930^ fails to co-localize with tubulin, but has staining at the apical cap and basal end of the parasite (arrowheads); Magenta: rabbit anti-HA; Green: mouse anti α-tubulin. J) IFA of reveals IMT1^Δ2-930^ co-localizes with ISP1 at the apical cap of the parasite; Magenta: mouse anti-ISP1; Green: rabbit anti-HA. K) IMT1^Δ2-1115^ mislocalizes and primarily is observed in the region corresponding to the parasite’s nucleus. Magenta: mouse anti-HA; Green: rabbit anti-IMC6. All scale bars = 2 μm.

### IMT1 contains a C-terminal domain that is sufficient for binding to the IMC

Our C-terminal and N-terminal deletion analyses pointed towards residues 930-1115 as being critical for IMT1 peripheral localization. To determine if this region was sufficient for localization to the IMC, we expressed just this portion of the protein (IMT1^930-1115^) in wildtype parasites (Fig 5A). IFA analysis showed bright staining at the periphery of the parasite that colocalized with IMC6, demonstrating that this region is indeed sufficient to bind to the IMC (Fig 5B). As expected, parasites expressing IMT1^930-1115^ did not localize to the SPMTs (Fig 5B, C). We therefore named this region of the protein the IMT1 IMC- binding region (IMT1^IBR^). To determine whether the IMT1^IBR^ was tethered to the IMC network or only the IMC membrane, we performed a detergent fractionation as described above. The IMT1^IBR^ partitioned largely to the insoluble cytoskeletal fraction indicating this portion of the protein is firmly attached to the IMC cytoskeleton and cannot be released by detergent extraction (Fig 5D).

**Figure 5.**
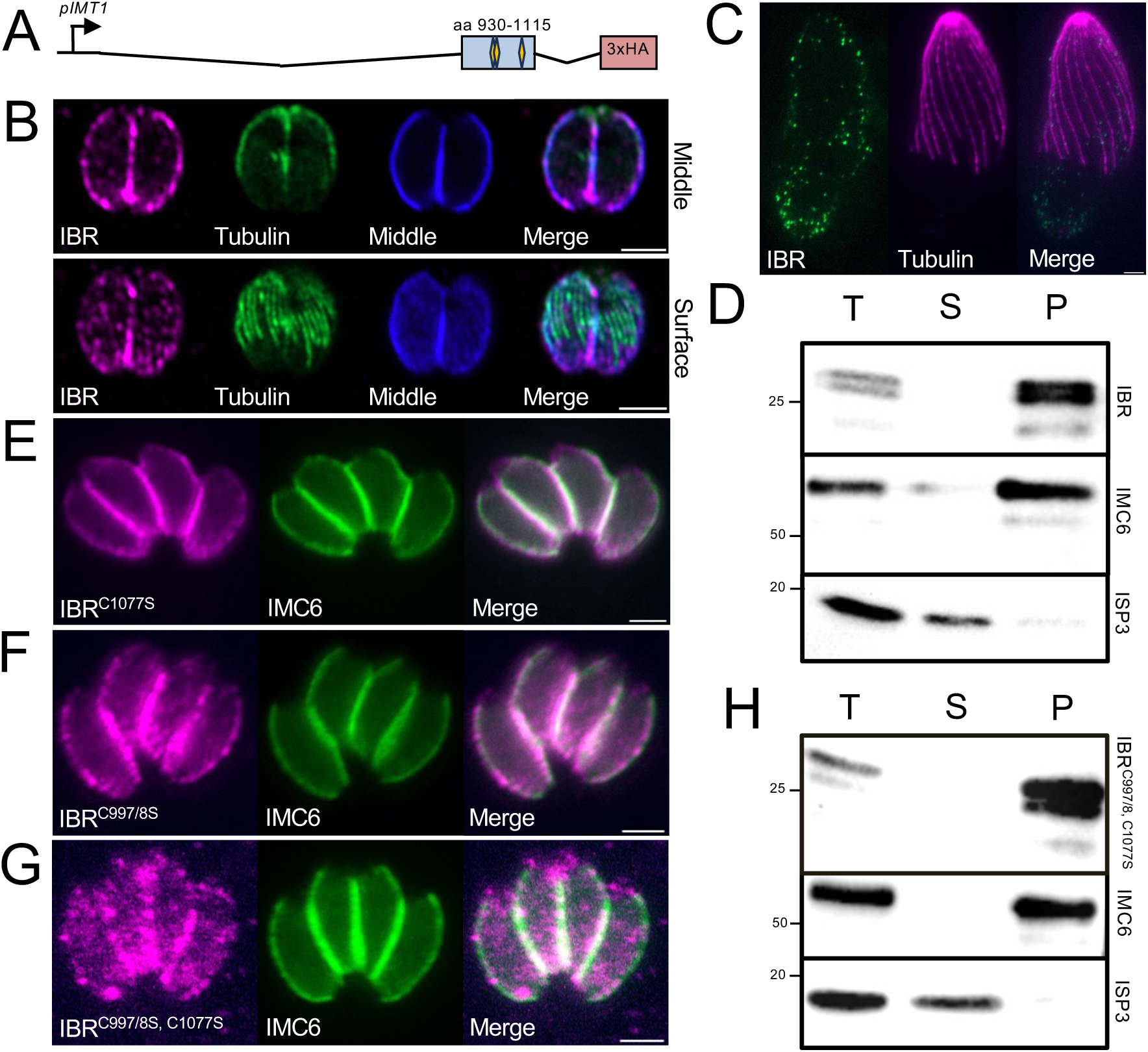
IMT1^Δ2-930^ is sufficient for IMC localization and is firmly associated with the IMC network. A) Gene model of IMT1^Δ930-1115^ driven from the endogenous promoter and fused to a 3xHA-epitope. B) IFA showing the IMT1^Δ930-1115^ is sufficient for localization to the IMC but not for localization to the SPMT. This region is denoted IMT1^IBR^ for IMT1 IMC binding region; Magenta: mouse anti-HA; Green: eGFP-Tubulin; Blue: rabbit anti- IMC6. C) U-ExM shows that the IMT1^IBR^ fails to target to the SPMTs. Green: rabbit anti- HA; Magenta: mouse anti α-tubulin. D) Detergent fractionation showing the IBR is still tethered to the cytoskeletal fraction. IMC6 and ISP3 serve as cytoskeletal and membrane fraction controls respectively. E-G) Mutagenesis of three putative palmitoylation sites in the IMT1^IBR^ reveals the mutation of C997S and C998S and the mutation of the third site C1007S individually do not affect IMT1^IBR^ localization. However, the mutation of all three sites causes a spotted localization at the periphery and dispersed signal in the body of the parasite; Magenta: mouse anti-HA; Green: rabbit anti-IMC6. H) Detergent fraction of IMT1^IBR(C997/998, C1077S)^ shows that mutagenesis of predicted palmitoylation sites does not affect IMT1^IBR^ tethering to the cytoskeleton despite the partial mislocalization. All scale bars = 2 μm.

CSS-Palm 4.0 weakly predicts three palmitoylation sites at residues 997, 998, and 1077 within the IMT1^IBR^ (31). To assess the role of palmitoylation in IMT1^IBR^ localization, we constructed a double mutant of residues 997 and 998 as well as a single mutant of residue 1077 by mutating the cysteines to serine. Both IMT1^IBR(C997/8S)^ and IMT1^IBR(C1077S)^ colocalized well with IMC6 at the periphery, matching the localization of the IMT1^IBR^ construct (Fig 5E, F). However, the mutation of all three putative palmitoylation sites (IMT1^IBR (C997/8S, C1077S)^) resulted more spotted IMC signal at the periphery of the parasite (Fig 5G). To assess if the IMT1^IBR (C997/8S, C1077S)^ mutant was still tethered to IMC network, we performed detergent fractionation on IMT1^IBR (C997/8S, C1077S^ and found it still partitioned to the cytoskeletal fraction (Fig 5H). This data indicates that while localization is partially affected in the triple mutant, incorporation into the IMC cytoskeletal network is unaffected.

### Proximity labeling identifies candidate IMT1 binding partners

To investigate which proteins mediate IMT1 binding to either the IMC or SPMTs, we performed proximity biotinylation using IMT1 as the bait protein (24, 41). To do this, we created a fusion of IMT1 with BioID^3xHA^ using C-terminal endogenous gene tagging (Fig. 6A). IFA analyses showed that the fusion protein targeted correctly to the IMC and SPMTs and that it biotinylates targets in these locations upon addition of biotin to the media (Fig. 6B). We have previously shown that detergent fractionation of cytoskeletal targets removes much of the background from proximity labelling experiments, so we employed this preparation for our large-scale biotinylation experiment (5). The biotinylated proteins in the cytoskeletal fraction were isolated by streptavidin chromatography and identified by tryptic digestion and mass spectrometry. The identified proteins were ranked by average NSAF score across two trials after deducting the control sample score and excluding any proteins that did not meet >2-fold enrichment. The top BioID hits identified were comprised of largely IMC and microtubule-associated proteins, consistent with the localization of IMT1 (Fig 6C, Table S1). This includes the IMT1 bait protein, an array of IMC proteins, the microtubule components alpha and beta tubulin, and the MAP TLAP2 in the top 30 proteins.

**Figure 6.**
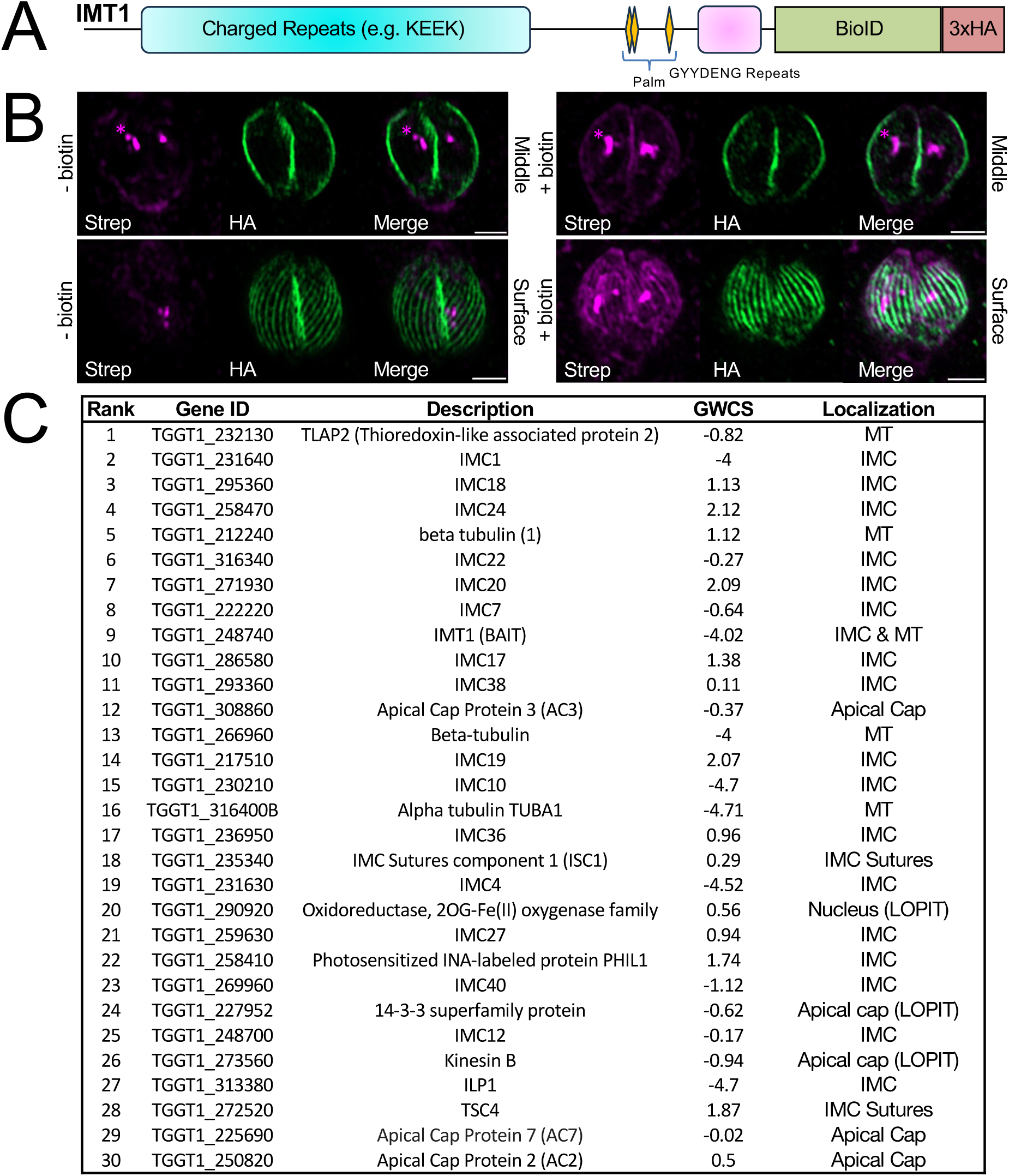
Proximity labelling using IMT1 as bait identifies candidate interacting partners. A) Gene model of the endogenous C-terminal tagging of IMT1 with a BioID^3xHA^ tag. B) IFA showing that the IMT1^BioID^ fusion targets to the IMC and SPMT similar to the wildtype protein and is active as assessed by streptavidin staining upon addition of biotin to the media. Asterisks note the endogenously biotinylated signal in the apicoplast. Minus and plus biotin are shown for both middle and surface views. Magenta: streptavidin-496; Green: mouse anti-HA. Scale bars = 2 μm. C) Table of the top 30 hits from streptavidin purification from IMT1^BioID^ lysates from the cytoskeletal fraction. Candidate proteins were ranked by average NSAF scores across two replicates following subtraction of control samples generated in parallel using untagged parasites plus biotin. GWCS = phenotype scored in the genome-wide CRISPR/Cas9 screen.

### IMT1 plays a role in the maintenance of TLAP2 levels on the SPMTs

The top hit in our IMT1 proximity labeling experiment was the microtubule associated protein TLAP2 (Fig. 6C) (22). To explore the relationship between IMT1 and TLAP2, we first colocalized TLAP2 and IMT1 and found they overlapped on the microtubules in maternal parasites, with TLAP2 also present in daughter buds as previously reported (Fig 7A, B) (22). We then endogenously tagged TLAP2 in the IMT1^3xHA^, Δ*imt1,* and IMT1^comp^ strains. We found that TLAP2 was present on the SPMTs in the Δ*imt1* strain, but the staining was clearly weaker than in the IMT1^3xHA^ line, suggesting a substantial decrease in protein levels in the IMT1 knockout (Fig 7C, D). We also observed that the staining along the SPMTs frequently did not extend as far down the microtubules, often terminating approximately halfway down the length of the parasite (Fig 7C, D, insets). The TLAP signal was restored in the IMT1^comp^ strain, demonstrating that the effect was due to the presence of IMT1 (Fig 7E). We used western blot analysis to quantify the loss of TLAP2 and found it was 72.3% reduced in Δ*imt1* parasites (Fig 7F, G). To address whether the TLAP2 similarly affects IMT1, we also disrupted *TLAP2* in IMT1^3xHA^ parasites and found no difference in the localization or levels of IMT1 (Fig S4). Together, this data shows that IMT1 plays a role in maintaining proper levels of TLAP2 on the SPMTs.

**Figure 7.**
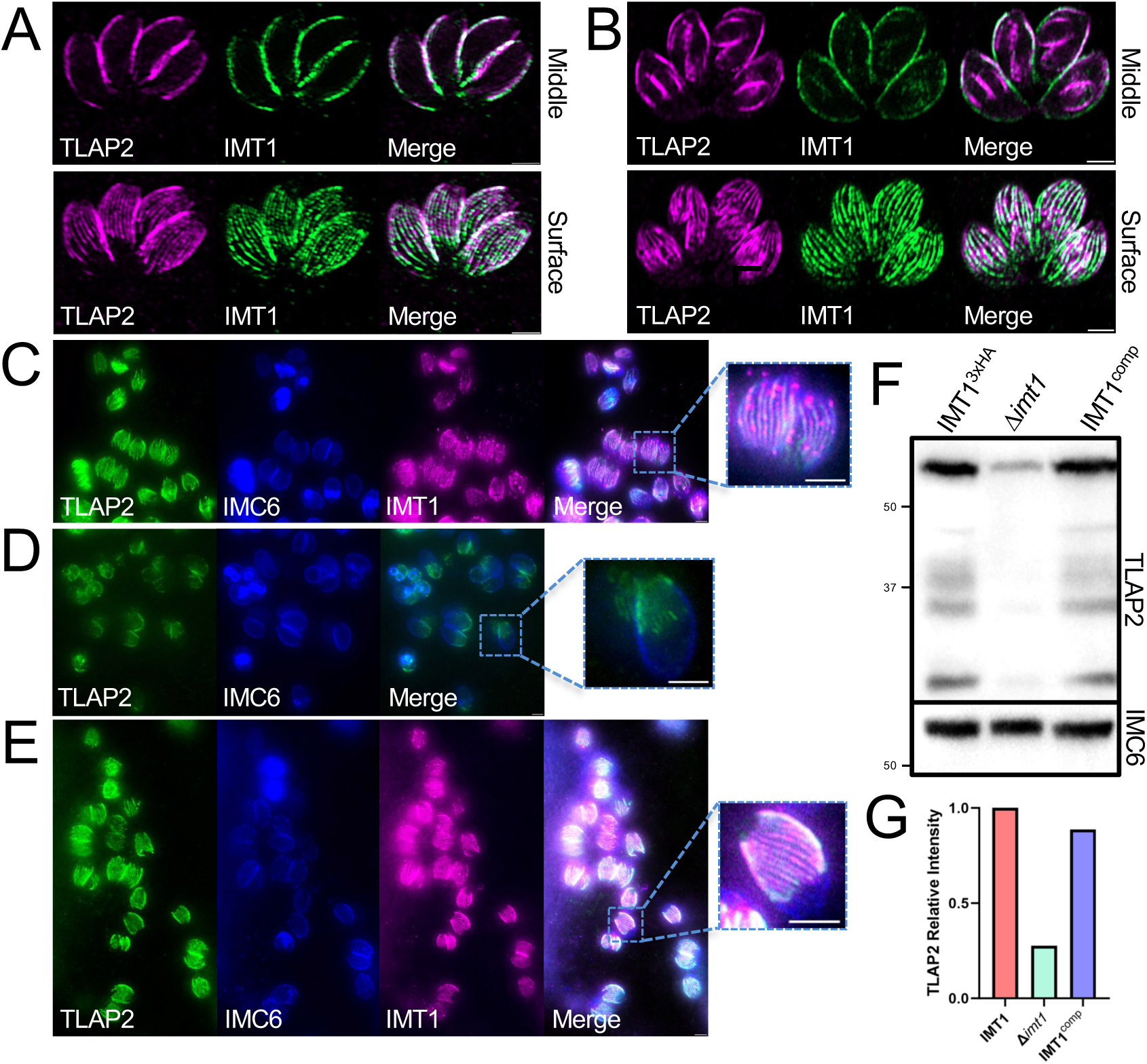
Disruption of IMT1 dramatically effects TLAP2 levels on the SPMTs. A) IFA showing that TLAP2^2xStrep3xTy^ and IMT1^3xHA^ colocalize in both middle and surface planes of maternal parasites. Magenta: mouse anti-Ty; Green: rabbit anti-HA. B) IFA of TLAP2^2xStrep3xTy^ showing localization to daughter buds in parasites undergoing endodyogeny, but IMT1^3xHA^ is restricted to the maternal parasite; Magenta: mouse anti-Ty; Green: rabbit anti-HA. C) Field of TLAP2^2xStrep3xTy^ IMT1^3xHA^ wildtype parasites showing TLAP2 co-localizes with IMT1 and extends down the length of the SPMTs. Inset shows a single vacuole with typical SPMT staining; Green: mouse anti-Ty; Blue: rat anti-IMC6; Magenta: rabbit anti-HA. D) Field of TLAP2^2xStrep3xTy^ tagged in *Δimt1* parasites showing TLAP2 signal is diminished. Inset shows a single vacuole with staining on the SPMTs extending less that in the control strain; Green: mouse anti-HA; Blue: rat anti-IMC6. E) Field of TLAP2^2xStrep3xTy^, IMT1^3xHA^ complemented parasites showing TLAP2 length and intensity is restored. Inset shows a single vacuole with rescued SPMT staining; Green: mouse anti- Ty; Blue: rat anti-IMC6; Magenta: rabbit anti-HA. F) Western blot of IMT1^3xHA^, Δ*imt1* and IMT1^comp^ parasites shows comparable expression levels of TLAP2 protein in HA-tagged and complemented stains, but a reduction of TLAP2-staining in Δ*imt1* parasites. IMC6 is used as a load control. G) Quantification of TLAP2 protein levels using Image Lab densitometry performed relative to IMC6 levels reveals dramatic 72.3% decrease in TLAP2 expression in the Δ*imt1* strain, which is restored by complementation. All scale bars = 2 μm.

### IMT1 directly binds to IMC1 via the IMT1^IBR^

The second highest ranking hit from our proximity labelling experiments was the and the IMC cytoskeletal protein IMC1 (42). We were interested in IMC1 as it is an integral component of the IMC cytoskeleton and is predominantly in the maternal IMC (Fig. 8A). Colocalization of IMC1 and IMT1 showed considerable overlap in maternal parasites, although smaller amounts of IMC1 are also present in the apical cap has been seen previously (5, 13). To determine if the IMT1^IBR^ and IMC1 directly bind one another, we assessed their interaction via pairwise Y2H assays (26). To do this, we generated Y2H bait and prey constructs for the IMT1^IBR^ and residues 14-591 of IMC1, which lack its potential palmitoylation sites (Fig 8B). We then co-transfected the constructs into the appropriate yeast strains and spotted them onto permissive and restrictive media. The spot assays showed a clear interaction of IMT1^IBR^ and IMC1 on restrictive media, demonstrating that these proteins do indeed interact (Fig 8C). This data demonstrates that IMT1 is tethered to the IMC at least in part via direct binding to the alveolin IMC1.

**Figure 8.**
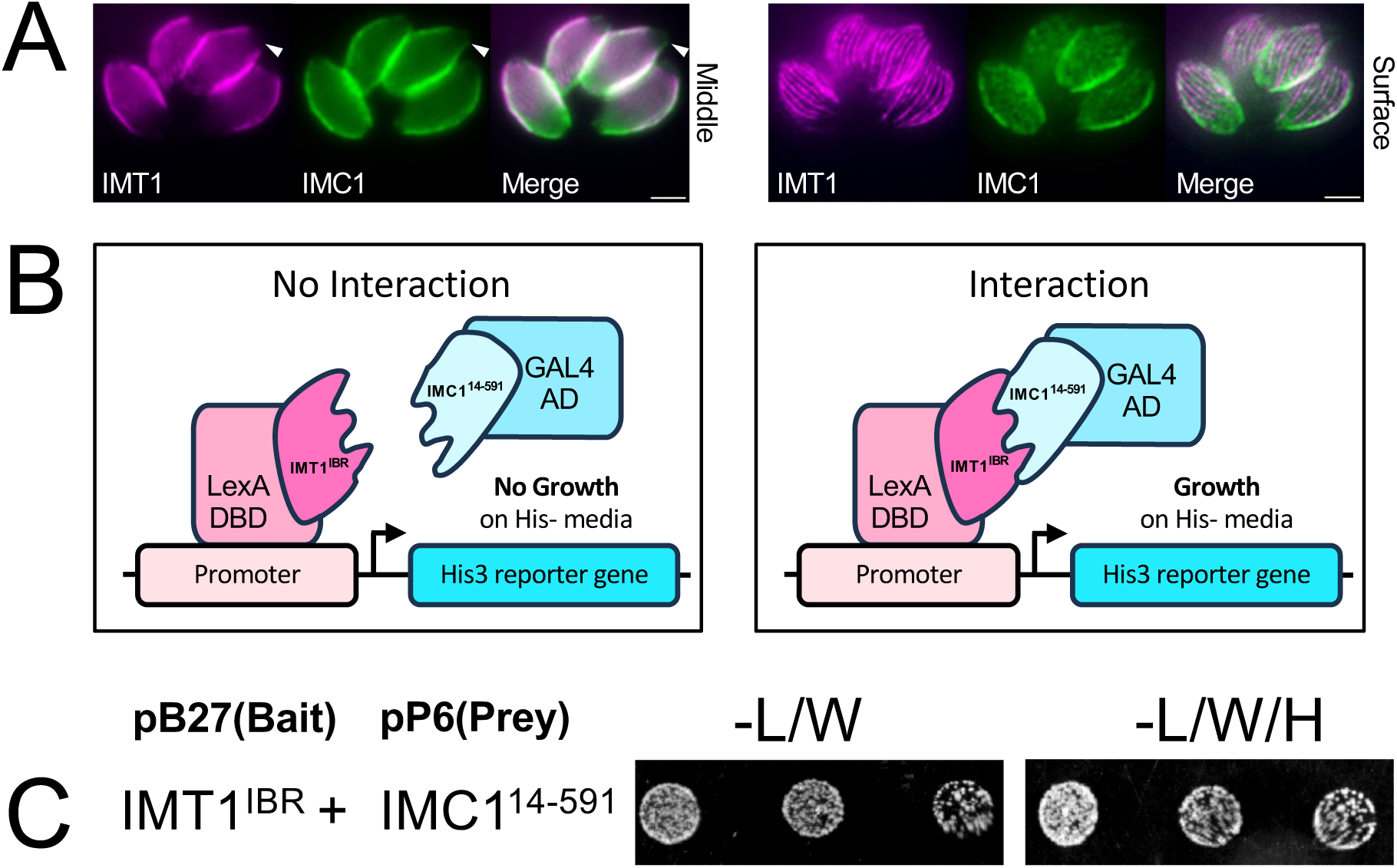
IMT1^IBR^ directly binds to IMC1 in the IMC network. A) IMT1^3xHA^ co-localized with IMC1 in middle and surface planes shows colocalization in the IMC, but IMC1 extends beyond into the apical cap (arrowheads); Magenta: rabbit anti-HA; Green: mouse anti-IMC1. Scale bars = 2 μm B) Diagram of non-interacting and interacting conditions of Y2H assay. C) Pairwise Y2H analysis of the IMT1^IBR^ and IMC1 shows growth on permissive media (-L/W) for both and restrictive media (-L/W/H) for IMC1, demonstrating that IMC1 is a direct-binding partner of IMT1 via the IMT1^IBR^.

## Discussion

In this work, we characterize the protein IMT1 which bridges the SPMTs and the IMC at the periphery of the parasite. The first clue that IMT1 differed from typical MAPs was that it localizes to both the SPMTs and the IMC body but is absent from the apical cap region of both of these cytoskeletal structures. Interestingly, TLAP2 is similarly absent from most of the apical cap region but also localizes to a small area below the apical polar ring at the extreme apex of the parasite (22). SPM2 also shows a differential localization in that it is absent from both the apical and basal portion of the SPMTs and thus is restricted to a central portion of the filaments (21). Like other SPMT associated proteins, SPM2 and TLAP2 also localize to the microtubules of developing daughter buds whereas IMT1 is limited to the maternal IMC and SPMTs, suggesting IMT1’s role is restricted to bridging the IMC and SPMTs once replication has concluded (21, 22). As it is difficult to distinguish the SPMTs from the IMC at the periphery of the parasite when viewing the IMC in the central plane of the parasite, it is possible that some of the other MAPs contain connections to the IMC that have not been previously identified.

Given the strongly negative phenotype score assigned to IMT1 in the Toxoplasma GWCS, it is surprising that we did not observe any fitness defect upon disruption of its gene (37). It is possible that this is the result of compensation upon gene knockout as compensation has been identified in other circumstances in *T. gondii* (34, 43, 44). The one protein we identified with poor similarity to IMT1 appears to be undetectable in tachyzoites whether or not IMT1 is present, suggesting that if compensation is occurring, it is via other means. To further address this, we attempted to assess function using the auxin-inducible degron system, but found that the knockdown appeared to be incomplete and we could not observe any phenotype in the knockdown parasites. We also attempted a conditional knockdown using promoter replacement with a tetracycline-regulatable promoter, but the approach failed as no regulation was observed. Disruption of *IMT1* in type II parasites again supports the lack of a phenotype in vitro or in vivo upon loss of the protein. It is also possible that the phenotype score from the initial GWCS is not reflective of the fitness defect of the loss of IMT1, highlighting the importance of interrogating each gene individually (37). Together, this data suggests that IMT1 either undergoes rapid compensation upon its absence or is dispensable under the conditions and life cycle stages evaluated here. It will be interesting to determine if other proteins that lack similarity to IMT1 similarly bridge the IMC and SPMTs and enable the close association of these peripheral cytoskeletal compartments.

Our deletion analyses identified regions IMT1 that are necessary for binding the SPMTs and the IMC. Of particular interest was the IMT1^IBR^ which is both necessary and sufficient for binding to the IMC. Mutagenesis of the three poorly predicted palmitoylation sites affected, but did not completely block, targeting to the IMC. This questions whether these sites are truly palmitoylated or if the triple cysteine mutation disrupted the structure of the protein fragment, which would best be resolved by metabolic radiolabeling studies (45). Regardless, IMT1^IBR^ is also firmly anchored into the cytoskeletal network of the IMC as shown by detergent extraction, indicating linkage to the IMC network. Our deletion analyses also demonstrated that the N-terminal charged region of IMT1 is necessary for binding to the SPMTs. However, this region does not appear to be sufficient for binding the SPMTs without the IMT1^IBR^. This suggests that incorporation of the protein into the IMC might be required prior to SPMT binding, although it is possible that deletion of the IMT1^IBR^ results in misfolding of the N-terminal binding region that blocks binding to the SPMTs. More refined deletion and mutagenesis experiments are likely to define the precise determinants that are both necessary and sufficient for binding the IMC and SPMTs.

We found that proximity labelling proved to be extremely useful for identifying candidate interactors of IMT1. As we have previously observed for the cytoskeletal IMC suture protein ISC4, the initial cytoskeletal fractionation of the BioID samples of a cytoskeletal fraction by detergent extraction results in the dramatic reduction of background proteins (5). This approach enabled us to hone in on TLAP2 and IMC1 as the top two candidate IMT1 interactors. Our Y2H analyses verified a direct interaction of IMC1 with the IMT1^IBR^ in the IMC, suggesting that this is how IMT1 is anchored into the maternal IMC cytoskeleton. However, while IMC1 is most robustly detected in the IMC body, it also extends into the apical cap, which lacks IMT1. This could suggest that IMT1 also interacts with other top hits such as IMC18 and IMC24, which are both restricted to the maternal IMC cytoskeleton and are absent from the apical cap (5). Both IMC18 and IMC24 are predicted to be dispensable (5, 37), and their functions have not been explored, thus gene knockout and two-hybrid analyses will be reveal if these proteins represent additional IMC linkages that restrict IMT1 from the apical cap.

We also identified TLAP2 as a candidate binding partner on the SPMTs via proximity labelling and found that TLAP2 is dramatically reduced on the SPMTs in Δ*imt1* parasites. This suggests a direct interaction between the proteins but could also be the result of IMT1 binding to other factors on the SPMTs or the SPMTs themselves. Alpha and beta tubulin were also high ranking in our dataset, which suggests that IMT1 directly binds SPMTs, where it aids in TLAP2 binding. We attempted to express full length IMT1 in human cells but were unable to see expression of the protein, perhaps due to its unusual sequence. In conclusion, this study identifies a link between the IMC and SPMTs of *T. gondii* which aids in the organization of TLAP2 on the SPMTs and may serve as a bridge between the two structures at the periphery of the parasite.

## Materials and Methods

### Toxoplasma and host cell culture

1. *T. gondii* RHΔ*ku80*Δ*hxgprt*, PruΔ*ku80*Δ*hxgprt*, and modified strains were cultured on confluent monolayers of human foreskin fibroblast (HFF) host cells at 37°C and 5% CO2 in Dulbecco’s Modified Eagle Medium (DMEM) supplemented with 10% fetal bovine serum, as previously described (38). Positive clonal selection for selectable markers was performed using 1 µM pyrimethamine, 50 µg/mL mycophenolic acid/xanthine, and 1 µM chloramphenicol (26). Homologous recombination at the UPRT locus was negatively selected for using 5 µM 5-fluorodeoxyuridine (FUDR) (38).

### Antibodies

Western and immunofluorescence assays (IFA) were probed using the following antibodies: mouse monoclonal antibody (mAb) HA.11 (Biolegend; 901515), rabbit polyclonal antibody (pAb) anti-HA (Invitrogen; P1715500), rat anti-HA (Roche; 3F12), rabbit pAb anti-IMC6 (46), rat pAb anti-IMC6 (46), mouse anti-ISP3 (34), mouse anti-α tubulin (Millipore Sigma; T6074), mouse anti-IMC1 (47), and mouse anti-ISP1 (34) . Biotinylated proteins were detected with streptavidin Alex Fluor 594 conjugate (Invitrogen; S-11227)

### Immunofluorescence assays (IFA) and Western blot

For IFA, epitope-tagged parasites were allowed to infect a layer of HFFs grown to confluence on glass coverslips. Coverslips were fixed with 3.7% formaldehyde, permeabilized with 3% BSA/PBS/0.2% Triton X-100, probed with primary antibodies, washed in 1XPBS, and probed with secondary antibodies conjugated to Alexa 594/488/504. Coverslips were mounted with Vectashield (Vector Labs) and fluorescence was captured using an Axio Imager.Z1 fluorescence microscope or a Zeiss LSM 980 with Airyscan 2. Images were processed with Zen software (Zeiss).

For Western Blotting, extracellular parasites were pelleted and washed with 1X PBS before lysis in Laemmli sample buffer and 100 mM DTT. Lysates were boiled at 100°C for 10 min and resolved by SDS-PAGE before transfer onto nitrocellulose membranes. Blots were then probed with primary antibodies, followed by secondary antibodies conjugated to horseradish peroxidase (HRP). Target proteins were visualized by chemiluminescence induced by Supersignal West Pico substrate (Pierce). Quantification of Western Blots was performed via BioRad Image Lab software normalized to load control (IMC6) and relative expression levels were calculated from volume normalized to the wild-type control.

### Ultrastructure-expanded Microscopy (U-ExM)

Extracellular parasites were fixed in 3.7% formaldehyde and added onto Poly-D-Lysine (A3890401, Gibco) coated coverslips, which were then incubated overnight at 37°C in a blocking fixative of 1.4% formaldehyde /2% acrylamide to facilitate cross-linking. A 19% sodium acrylate/10% acrylamide/0.1% BIS gel was formed atop the coverslip, cured for 1 hour at 37°C, separated, and denatured for 1.5 h at 95 °C. The gel was then expanded in deionized water, measured, then shrunk in PBS for primary antibody and secondary antibody incubations at RT, overnight and for 2.5 hours respectively. The final gels were cut and mounted onto Poly-D-Lysine coated coverslips and viewed via fluorescence microscopy as described above (35, 48).

### Epitope tagging and Gene Disruption

Proteins were endogenously tagged at the C-terminus using CRISPR/Cas9. A sgRNA 100-200 base pairs downstream of the stop codon was ligated into a pU6-Universal plasmid, as previously described (26, 49). Homology-directed repairs (HDR) templates were amplified from LIC (ligation-independent cloning) vectors including epitope tags (e.g. 3xHA, 2xStrep3xTy, smHA) and selectable markers via primers with 40 bp of homology upstream and downstream of the sgRNA. Primers P1-P12 were used for pU6- Universal plasmid and HDR templates (Table S2).

For gene knockout, a sgRNA was designed to target the coding region of the gene interest and ligated into a pU6-Universal plasmid (49). HDR templates were amplified from a pJET vector containing the HXGPRT selectable marker driven by the NcGRA7 promoter using primers with homology to 40 bp upstream of the start codon and downstream of the region of integration from endogenous tagging. *IMT1* and *TLAP2* were knocked with primers P15-P19 and P24-P27 respectively (Table S2). Approximately 50µg of modified pU6-Universal plasmid was electroporated with 400 µL of phenol- chloroformed and ethanol precipitated PCR-amplified HDR template into the appropriate parasite strains for integration of epitope tagging and knockout constructs. Transfected lines were allowed to infect confluent HFF monolayers before the appropriate drug selection was applied. Successful tagging and selection were confirmed via IFA before limiting dilution was performed to obtain clonal lines. Knockouts were additionally verified via PCR amplification of the HXGPRT caseate and failure to amplify the coding region. P20-P23 and P28-P31 were used to verify knockouts (Table S2).

### Plaque Assay

Parasites were allowed to infect 6-well plates seeded with a confluent monolayer of HFFs and undergo repeated iterations of the lytic cycle for 7 days (RHΔ*ku80*Δ*hxgprt*) or 9 days (Pru Δ*ku80*Δ*hxgprt*). The plates were washed with PBS and fixed with 100% ice cold methanol for 5 min. before staining with crystal violet. The area of ∼50 plaques per condition were measured using Zen software (Zeiss). All conditions were performed in biological replicates and statistical significance was calculated using unpaired t-tests.

### Mouse Brain Cyst Quantification

Wildtype PruΔ*ku80*Δ*hxgprt* and Δ*imt1II* parasites were resuspended in Opti-MEM and 300 of each were injected intraperitoneally into groups of female CBA/J mice. Parasite viability was confirmed via plaque assay. Mice condition and weight were monitored over the course of 30 days post-injection. The mice were euthanized as per AVMA guidelines— using slow (20–30% per minute) displacement of chamber air with compressed CO2 followed by cervical dislocation. Mouse brains were collected and weighed before homogenization for enumeration of LDH2-GFP+ tissue cysts. Equivalent proportions of each brain were screened by fluorescence microscopy for cyst enumeration.

### Detergent extractions

For detergent extraction to separate the parasite’s cytoskeleton from soluble material, extracellular parasites were washed in 1X PBS, pelleted, and lysed in 1 mL of 1% Triton X-100 lysis buffer (50mM Tris-HCl [pH 7.4], 150mM NaCl) supplemented with Complete Protease Inhibitor Cocktail (Roche) for 20 min on ice. The lysates were centrifuged for 10 min at 16,000 x *g* at 4°C. Equivalent amounts of total, supernatant (detergent soluble), and pellet (detergent insoluble) fractions were separated by SDS-PAGE and analyzed by Western blot. IMC6 and ISP3 served as controls for the cytoskeletal and membrane fractions, respectively (34, 46).

### Deletion-Analysis and Complementation of *Δimt1* parasites

The endogenous promoter amplified from genomic DNA and the full-length coding region of IMT1 amplified from cDNA and genomic DNA in three fragments, was cloned into a UPRT targeting vector via Gibson Assembly using primers P32-P41 (Table S2). Palmitoylation site mutants and the N-terminal and C-terminal deletion series were generated from this full-length construct via Q5 site-directed mutagenesis using primers P42-P61 (Table S2).

All constructs were linearized via Psil-HFv2 and prepared and transfected as described above into Δ*imt1* parasites for complementation or WT parasites for deletion analyses. Selection with 5 μg/mL 5-fluorodeoxyuridine (FUDR) for replacement of UPRT locus was performed after 2 passages and clonal lines were obtained following IFA confirmation of integration and limiting dilution.

### Pairwise Yeast-Two Hybrid analyses

Regions of IMC1 and IMT1 were cloned via Gibson Assembly into the pB27 (N-LexA-bait-C fusion) vector as a N-terminal fusion of the LexA DNA binding domain or pP6 (N- GAL4AD-prey-C fusion) vector as a N-terminal fusions of the GAL4 Activation domain (Hybrigenics Services) using primers P62-P73 (Table S2). Potential binding partners were co-transformed into the L40 strain of *S. cerevisiae* [MATa his3D200trp1-901 leu2-3112 ade2 LYS2::(4lexAop-HIS3) URA3::(8lexAop-lacZ) GAL4] before plating onto permissive (-Leu/-Trp) plates to obtain clonal strains. These clones were expanded in permissive (-Leu/-Trp) media before dilution to an OD600 of 0.2, and spotting onto both permissive (-Leu/-Trp) and restrictive (-Leu/-Trp/-His) plates in a 2x dilution series. Interaction was assessed after 4-5 days of growth.

### Modification of eGFP expressing TgTubA1

Flanks with homology to the *Ku80* locus were amplified and cloned into the eGFP.TgTubA1.CAT plasmid via Gibson Assembly using primers p74-83 (Table S2). For expression of eGFP-tubulin, 100 μg of eGFP.TgTubA1.ku80.CAT was linearized via Not1-HF and transfected with sgRNA targeted to the *Ku80* locus and selected via 1 µM chloramphenicol. Clonal lines were obtained following selection using limiting dilution and confirmed via IFA.

### Affinity purification of biotinylated proteins

Parental and IMT1-BioID fusion parasites were allowed to infect and lyse confluent monolayers of HFF in media supplemented with 150 μM of biotin. Parasites were then pelleted and washed before lysis in RIPA buffer [1% Triton-X 100, 1% SDS, 150 mM NaCl, 50 mM Tris pH 7.4] supplemented with cOmplete Protease Inhibitor Mixture (Roche). After pelleting, the supernatant was discarded and the pellet containing the cytoskeletal fraction was incubated with Streptavidin High-Capacity Agarose (Pierce) overnight at 4°C. Beads were washed in RIPA and 8M urea before submission for mass spectrometry analysis. The samples were digested and proteins identified by mass spectrometry as described (28).

## Acknowledgements

We thank Gary Ward for IMC1 antibodies, Michael Reese for the eGFP-tubulin construct, and Silvia Moreno for the smHA construct. We also thank members of the Bradley lab for reading and editing of the manuscript. This work was supported by the Undergraduate Research Scholars Program (URSP) and a Whitcome Undergraduate Research Fellowship to E.S.C, as well as NIH grants #AI123360 to P.J.B. and #GM153408 to J.A.W. The funders had no role in the study design, data collection, interpretation, or decision to submit the work for publication.

---

**Figure S1.**
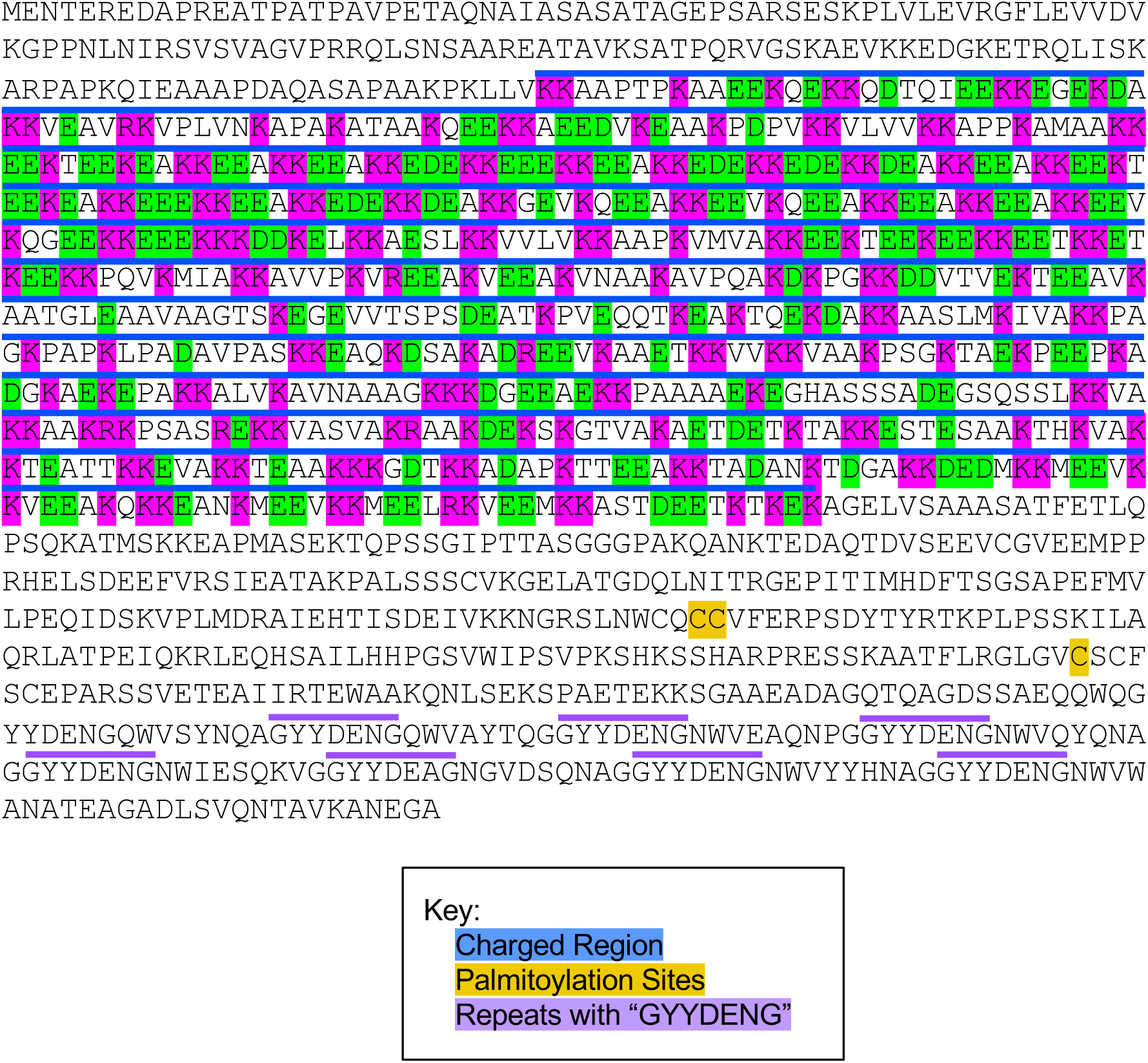
TGGT1_248740 sequence characteristics. Amino acid sequence of TGGT1_248740 highlighting its N-terminal charged region, three predicted palmitoylation sites, and C-terminal ∼15 amino acid repeats contain the sequence GYYDENG.

**Figure S2.**
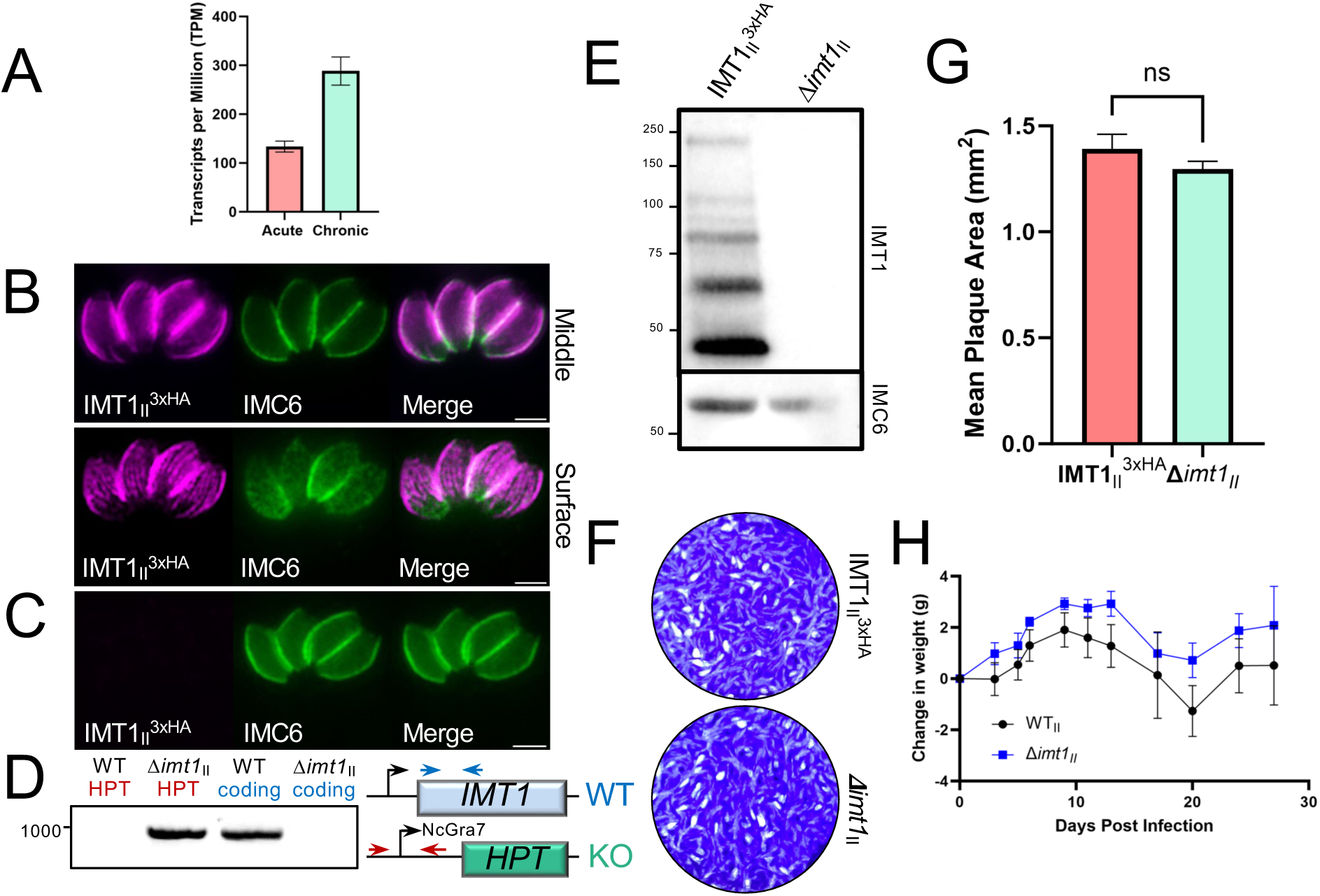
Gene knockout of *IMT1* in Pru Δ*ku80*Δ*hxgprt* parasites. A) Transcriptomic analysis reveals IMT1 is elevated in the chronic (28 dpi) compared to the acute (10 dpi) stage of infection. TPM = Transcripts per million (39). B) Endogenous 3xHA epitope-tagging of IMT1 in Pru Δ*ku80*Δ*hxgprt* parasites shows localization to the IMC (top panel: middle plane) and the SPMT (bottom panel: surface plane) which matches the localization seen in RH strain parasites (Fig 1). C) IFA of Δ*imt1II* parasites lacking staining for IMT1II ; Magenta: mouse anti-HA; Green: rabbit anti-IMC6. D) PCR and diagram showing amplification of the integrated selectable marker hypoxanthine-xanthine-guanine phosphoribosyl transferase (HPT) cassette but no amplification of the replaced IMT1 coding region. The converse is true for the wildtype control, and all bands match the sizes predicted by the primer positions shown in the diagram. E) Western blot of IMT1II and Δ*imt1II* strains confirms IMT1 staining for endogenously tagged, but not knockout parasites. F) Representative wells for IMT1^3xHA^ and Δ*imt1II* plaque assays 9 dpi. G) Quantification shows no significant difference in average plaque area between tagged and knockout strains. H) Graph showing change in CBA/J mice weights following intraperitoneal injection up until euthanizing and collection of brains for cyst enumeration. All scale bars = 2 μm.

**Figure S3.**
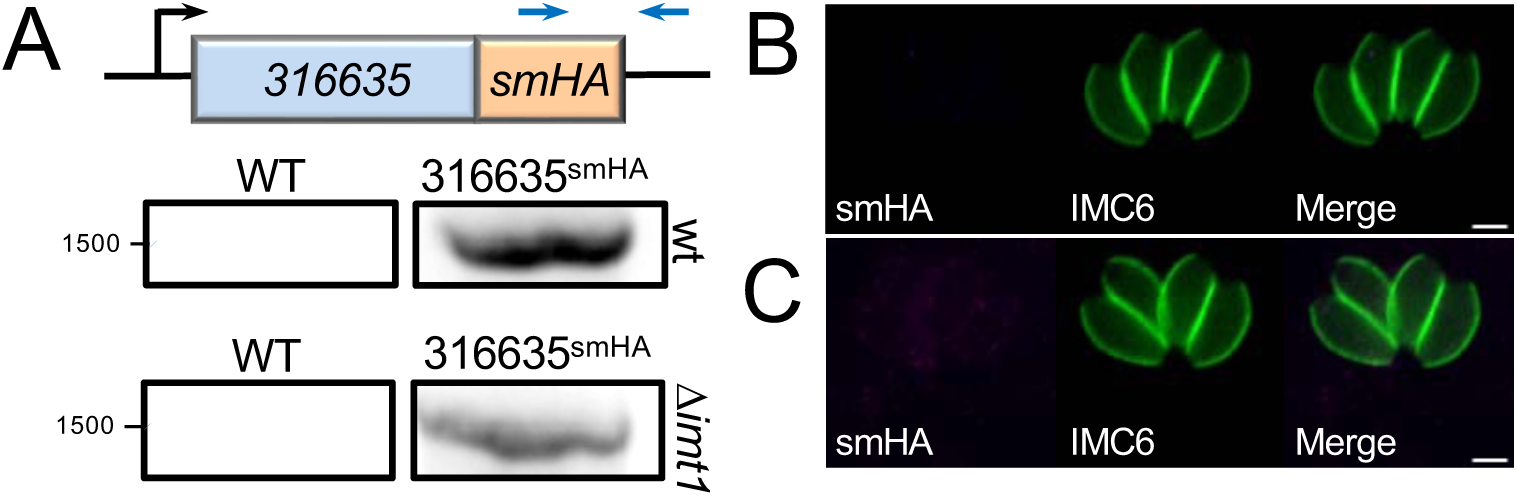
Endogenous gene tagging of TGGT1_316635 in wildtype and Δ*imt1* parasites. A) PCR and diagram confirming integration of the smHA-epitope tag at the C-terminus of TGGT1_316635. Amplicon for both TGGT1_316635^smHA^ and *Δimt1* TGGT1_316635^smHA^ strains matches the size predicted by the primer positions shown in the diagram. B) HA staining for TGGT1_316635^smHA^ fails to the reveal any staining suggesting that TGGT1_316635 is not expressed sufficiently to be detected; Magenta: mouse anti-HA; Green: rabbit anti-IMC6. C) Staining for TGGT1_316635^smHA^ in a *Δimt1* background also fails to show any detectable protein; Magenta: mouse anti-HA; Green: rabbit anti-IMC6. All scale bars = 2 μm.

**Figure S4.**
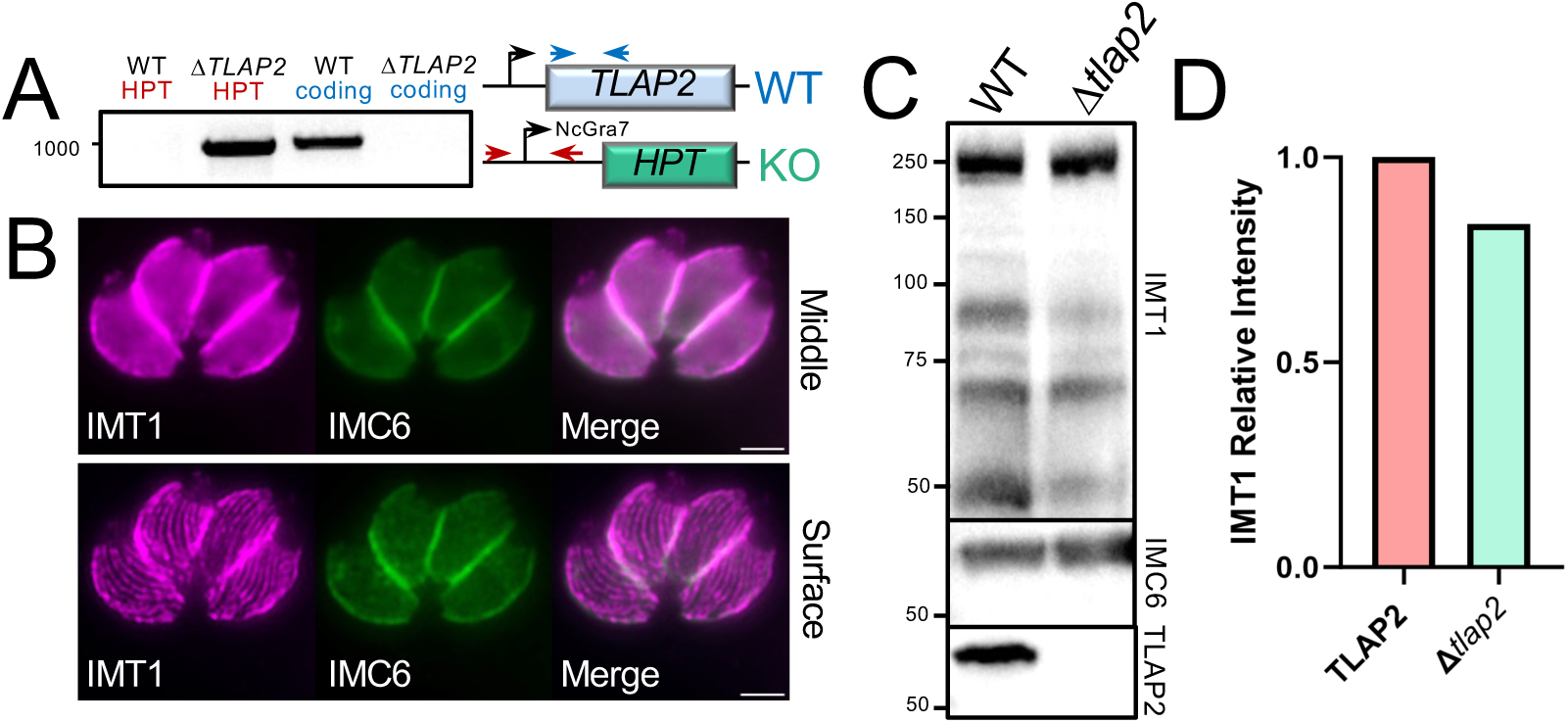
Disruption of TLAP2 does not have any significant effect on IMT1 localization or expression. A) PCR and diagram showing the Δ*tlap2* strain contains the (HPT) cassette and lacks the replaced TLAP2 coding region. The converse is true for the wildtype control, and all amplified bands match the sizes predicted by the primer positions shown in the diagram. B) IMT1^3xHA^ in a *Δtlap2* background showing localization to the IMC in a middle plane (top panel) and localization to the SPMT in a surface plane (bottom panel), matching wildtype localization; Magenta: mouse anti-HA; Green: rabbit anti-IMC6. Scale bars = 2 μm. C) Western blot of IMT1^3xHA^ and *Δtlap2* IMT1^3xHA^ showing comparable expression of IMT1 in both strains and the absence of TLAP2 expression in the knockout. D) Quantification of IMT1 protein levels using BioRad Image Lab densitometry performed relative to IMC6 levels shows comparable intensity between wildtype and *Δtlap2* strains.

**Table S1.Full dataset for IMT1 proximity labeling**. Full list of genes identified by IMT1^BioID^. NSAF scores are shown for each gene for the two replicates of untagged and BioID tagged sample plus biotin. Enrichment difference refers to the difference between the average NSAF score for the IMT1^BioID^ and control samples.

**Table S2.Oligonucleotide primers used in this study**. All primer sequences are shown in the 5’ to 3’ orientation.

